## Supplemental Materials for "The five homologous CiaR-controlled Ccn sRNAs of *Streptococcus pneumoniae* modulate Zn-resistance"

**Running Title:** Loss of the Ccn sRNAs cause Zn hypersensitivity

**Keywords:** small RNA; post-transcriptional regulation; Zn homeostasis; Mn homeostasis; CiaRH

### TABLE OF CONTENTS

**SUPPLEMENTAL TABLES**

**TABLE S1.** *S. pneumoniae* strains used in this study

**TABLE S2.** Primers used to construct mutants used in this study

**TABLE S3.** Comparison of gene expression *S. pneumoniae* D39 and derived Δ*ccnABCDE* strains in BHI broth by RNA-seq. (see Excel spreadsheet)

**TABLE S4.** Comparison of gene expression *S. pneumoniae* TIGR4 and derived Δ*ccnABCDE* strains in BHI broth by RNA-seq. (see Excel spreadsheet)

**TABLE S5.** Comparison of gene expression *S. pneumoniae* D39 and derived Δ*ccnABCDE* strains in BHI broth with 0.2 mM $ZnSO_4$ by RNA-seq. (see Excel spreadsheet)

**TABLE S6.** Oligonucleotide primers used for qRT-PCR

**SUPPLEMENTAL FIGURES**

**FIGURE S1.** Growth phenotypes of *S. pneumoniae* D39 derived strains harboring deletion of individual *ccn* genes.

**FIGURE S2.** Virulence phenotypes of *S. pneumoniae* strains harboring deletion of individual *ccn* genes.

**FIGURE S3.** Growth phenotypes of *S. pneumoniae* TIGR4 derived strains harboring deletion of the *ccn* genes.

**FIGURE S4.** Doubling times of *S. pneumoniae* D39, Δ*ccnABCDE* mutant, or Δ*ccnABCDE* mutant strain complemented with *ccnA, ccnB,* and *ccnC* in C medium alone or supplemented with Zn.

**FIGURE S5.** Growth phenotypes of *S. pneumoniae* D39 and derived strains harboring deletion of the *ccn* genes and/or *psaR* or *mntE*.

**SUPPLEMENTAL REFERENCES**

**SUPPLEMENTAL TABLES**

**Table S1.** *S. pneumoniae* strains used in this study

| Strain | Genotype (description)[a] | Antibiotic resistance[b] | Reference or source |
|---|---|---|---|
| K272 | D39 Δ*cps* Δ*pnp*::$P_c$-[*kan rpsL*$^+$] | $Kan^R$ | (*1*) |
| IU1781 | D39 *rpsL1* | $Str^R$ | (*2*) |
| IU5122 | D39 Δ*cps rpsL1* CEP::$P_c$-[*kan rpsL*$^+$] | $Kan^R$ | (*3*) |
| IU5382 | D39 Δ*cps* CEP::$P_{fcsK}$-*ftsE* | None | (*3*) |
| IU9681 | D39 Δ*cps hlpA hlpA-mkate-cat* | $Cm^R$ | Malcolm Winkler |
| IU10508 | D39 Δ*cps* Δ*bgaA*::*kan*-$T_1$-$T_2$-$P_{spd\_1874}$-*lacZ* | $Kan^R$ | (*4*) |
| IU11966 | TIGR4 | None | (*5, 6*) |
| IU12001 | TIGR4 Δ*cps* | None | (*7*) |
| NRD10068 | D39 *rpsL1* Δ*ccnA*::$P_c$-[*kan rpsL*$^+$] | $Kan^R$ | This study |
| NRD10069 | D39 *rpsL1* Δ*ccnB*::$P_c$-[*kan rpsL*$^+$] | $Kan^R$ | This study |
| NRD10070 | D39 *rpsL1* Δ*ccnC*::$P_c$-[*kan rpsL*$^+$] | $Kan^R$ | This study |
| NRD10071 | D39 *rpsL1* Δ*ccnD*::$P_c$-[*kan rpsL*$^+$] | $Kan^R$ | This study |
| NRD10072 | D39 *rpsL1* Δ*ccnE*::$P_c$-[*kan rpsL*$^+$] | $Kan^R$ | This study |
| NRD10073 | D39 *rpsL1* Δ*ccnA* | $Str^R$ | This study |
| NRD10074 | D39 *rpsL1* Δ*ccnB* | $Str^R$ | This study |
| NRD10075 | D39 *rpsL1* Δ*ccnC* | $Str^R$ | This study |
| NRD10076 | D39 *rpsL1* Δ*ccnD* | $Str^R$ | This study |
| NRD10077 | D39 *rpsL1* Δ*ccnE* | $Str^R$ | This study |
| NRD10078 | D39 *rpsL1* Δ*ccnB* Δ*ccnC*::$P_c$-[*kan rpsL*$^+$] | $Kan^R$ | This study |
| NRD10079 | D39 *rpsL1* Δ*ccnB* Δ*ccnC* | $Str^R$ | This study |
| NRD10080 | D39 *rpsL1* Δ*ccnB* Δ*ccnC* Δ*ccnD*::$P_c$-[*kan rpsL*$^+$] | $Kan^R$ | This study |
| NRD10081 | D39 *rpsL1* Δ*ccnB* Δ*ccnC* Δ*ccnD* | $Str^R$ | This study |
| NRD10083 | D39 *rpsL1* Δ*ccnB* Δ*ccnC* Δ*ccnD* Δ*ccnE*::$P_c$-[*kan rpsL*$^+$] | $Kan^R$ | This study |
| NRD10085 | D39 *rpsL1* Δ*ccnB* Δ*ccnC* Δ*ccnD* Δ*ccnE* | $Str^R$ | This study |
| NRD10171 | D39 *rpsL1* Δ*ccnAB*::$P_c$-[*kan rpsL*$^+$] Δ*ccnC* Δ*ccnD* Δ*ccnE* | $Kan^R$ | This study |
| NRD10176 | D39 *rpsL1* Δ*ccnAB* Δ*ccnC* Δ*ccnD* Δ*ccnE* | $Str^R$ | This study |
| NRD10220 | TIGR4 *rpsL1* | $Str^R$ | This study |
| NRD10225 | TIGR4 *rpsL1* Δ*ccnA*::$P_c$-[*kan rpsL*$^+$] | $Kan^R$ | This study |
| NRD10230 | TIGR4 *rpsL1* Δ*ccnA* | $Str^R$ | This study |
| NRD10247 | TIGR4 *rpsL1* Δ*ccnA* Δ*ccnB*::$P_c$-[*kan rpsL*$^+$] | $Kan^R$ | This study |
| NRD10249 | TIGR4 *rpsL1* Δ*ccnA* Δ*ccnB* | $Str^R$ | This study |
| NRD10251 | TIGR4 *rpsL1* Δ*ccnA* Δ*ccnB* Δ*ccnC*::$P_c$-[*kan rpsL*$^+$] | $Kan^R$ | This study |
| NRD10254 | TIGR4 *rpsL1* Δ*ccnA* Δ*ccnB* Δ*ccnC* | $Str^R$ | This study |
| NRD10257 | TIGR4 *rpsL1* Δ*ccnA* Δ*ccnB* Δ*ccnC* Δ*ccnD*::$P_c$-[*kan rpsL*$^+$] | $Kan^R$ | This study |
| NRD10261 | TIGR4 *rpsL1* Δ*ccnA* Δ*ccnB* Δ*ccnC* Δ*ccnD* | $Str^R$ | This study |
| NRD10265 | TIGR4 *rpsL1* Δ*ccnA* Δ*ccnB* Δ*ccnC* Δ*ccnD* Δ*ccnE*::$P_c$-[*kan rpsL*$^+$] | $Kan^R$ | This study |
| NRD10266 | TIGR4 *rpsL1* Δ*ccnA* Δ*ccnB* Δ*ccnC* Δ*ccnD* Δ*ccnE* | $Str^R$ | This study |

| | | | |
|---|---|---|---|
| NRD10306 | TIGR4 *rpsLK56T* Δ*ccnA*::P$_c$-[*kan rpsL*$^+$] | Kan$^R$ | This study |
| NRD10311 | TIGR4 *rpsL*$^+$*-rpsG*$^+$*-cat* | Cm$^R$ | This study |
| NRD10312 | TIGR4 *rpsLK56T* Δ*ccnA* | Str$^R$ | This study |
| NRD10322 | TIGR4 *rpsLK56T* Δ*ccnA* Δ*ccnB*::P$_c$-[*kan rpsL*$^+$] | Kan$^R$ | This study |
| NRD10324 | TIGR4 *rpsLK56T* Δ*ccnA* Δ*ccnB* | Str$^R$ | This study |
| NRD10330 | TIGR4 *rpsLK56T* Δ*ccnA* Δ*ccnB* Δ*ccnC*::P$_c$-[*kan rpsL*$^+$] | Kan$^R$ | This study |
| NRD10332 | TIGR4 *rpsLK56T* Δ*ccnA* Δ*ccnB* Δ*ccnC* | Str$^R$ | This study |
| NRD10336 | TIGR4 *rpsLK56T* Δ*ccnA* Δ*ccnB* Δ*ccnC* Δ*ccnD*::P$_c$-[*kan rpsL*$^+$] | Kan$^R$ | This study |
| NRD10340 | TIGR4 *rpsLK56T* Δ*ccnA* Δ*ccnB* Δ*ccnC* Δ*ccnD* | Str$^R$ | This study |
| NRD10344 | TIGR4 *rpsLK56T* Δ*ccnA* Δ*ccnB* Δ*ccnC* Δ*ccnD* Δ*ccnE*::P$_c$-[*kan rpsL*$^+$] | Kan$^R$ | This study |
| NRD10345 | TIGR4 *rpsLK56T* Δ*ccnA* Δ*ccnB* Δ*ccnC* Δ*ccnD* Δ*ccnE* | Str$^R$ | This study |
| NRD10346 | TIGR4 *rpsL*$^+$*-rpsG*$^+$*-cat* Δ*ccnA* Δ*ccnB* Δ*ccnC* Δ*ccnD* Δ*ccnE* | Cm$^R$ | This study |
| NRD10390 | D39 *rpsL1* CEP::P$_c$-[*kan rpsL*$^+$] | | This study |
| NRD10391 | D39 *rpsL1* Δ*ccnAB* Δ*ccnC* Δ*ccnD* Δ*ccnE* CEP::P$_c$-[*kan rpsL*$^+$] | Kan$^R$ | This study |
| NRD10393 | D39 *rpsL1* Δ*ccnAB* Δ*ccnC* Δ*ccnD* Δ*ccnE* CEP::T$_1$-T$_2$-*ccnA-ccnB* | Str$^R$ | This study |
| NRD10394 | D39 *rpsL1* Δ*ccnAB* Δ*ccnC* Δ*ccnD* Δ*ccnE* CEP::T$_1$-T$_2$-*ccnC* | Str$^R$ | This study |
| NRD10396 | D39 *rpsL1* Δ*ccnAB* Δ*ccnC* Δ*ccnD* Δ*ccnE* CEP::T$_1$-T$_2$-*ccnA-ccnB* Δ*bgaA*::*kan-T$_1$-T$_2$-ccnD* | Str$^R$, Kan$^R$ | This study |
| NRD10397 | D39 *rpsL1* Δ*ccnAB* Δ*ccnC* Δ*ccnD* Δ*ccnE* CEP::T$_1$-T$_2$-*ccnC* Δ*bgaA*::*kan-T$_1$-T$_2$-ccnD* | Str$^R$, Kan$^R$ | This study |
| NRD10441 | D39 *rpsL1* Δ*psaR*::P$_c$-[*kan rpsL*$^+$] | Kan$^R$ | This study |
| NRD10442 | D39 *rpsL1* Δ*mntE*::P$_c$-[*kan rpsL*$^+$] | Kan$^R$ | This study |
| NRD10443 | D39 *rpsL1* Δ*ccnAB* Δ*ccnC* Δ*ccnD* Δ*ccnE* Δ*psaR*::P$_c$-[*kan rpsL*$^+$] | Kan$^R$ | This study |
| NRD10444 | D39 *rpsL1* Δ*ccnAB* Δ*ccnC* Δ*ccnD* Δ*ccnE* Δ*mntE*::P$_c$-[*kan rpsL*$^+$] | Kan$^R$ | This study |
| NRD10447 | D39 *rpsL1* Δ*psaR* | Str$^R$ | This study |
| NRD10448 | D39 *rpsL1* Δ*mntE* | Str$^R$ | This study |
| NRD10449 | D39 *rpsL1* Δ*ccnAB* Δ*ccnC* Δ*ccnD* Δ*ccnE* Δ*psaR* | Str$^R$ | This study |
| NRD10450 | D39 *rpsL1* Δ*ccnAB* Δ*ccnC* Δ*ccnD* Δ*ccnE* Δ*mntE* | Str$^R$ | This study |
| NRD10533 | D39 *rpsL1* Δ*sodA::erm* | Erm$^R$ | This study |
| NRD10534 | D39 *rpsL1* Δ*ccnAB* Δ*ccnC* Δ*ccnD* Δ*ccnE* Δ*sodA::erm* | Erm$^R$ | This study |
| TIGR4S | TIGR4 *rpsLK56T* | Str$^R$ | This study |
| TIGR4SΔcps | TIGR4 *rpsLK56T* Δ*cps* | Str$^R$ | This study |

[a] Strains were constructed by transformation of amplicons into the indicated recipient strain as described in Materials and Methods. Primers used to synthesize fusion amplicons are listed in Table S2.

[b]Antibiotic resistance markers: Kan, kanamycin; Str, streptomycin; Erm, Erythromycin.

**Table S2.** Primers used to construct mutants used in this study

| Primer name | Primer Sequence (5'-3') | Template | Product |
|---|---|---|---|
| **For construction of strain NRD10068 (Δ*ccnA::*$P_c$-[kanR-rpsL$^+$])** | | | |
| Spn001 | TCATAGACAAGGCGACTGGTAAGG | IU1945 | Upstream of *ccnA* |
| Spn002 | CCATTAAAAATCAAACGGATCCTAAAAAAAGTTTAGGATTTTATTAAATAAAGTTAGG | | |
| kanrpsL For | TAGGATCCGTTTGATTTTTAATGGATAATG | K272 | $P_c$-[*kan rpsL*$^+$][b] |
| kanrpsL rev | GGGCCCCTTTCCTTATGCTTTTG | | |
| Spn003 | TCCAAAAGCATAAGGAAAGGGGCCCGGCTTTTTGCGTGGTGAGGTGCTGGTG | IU1945 | Downstream of *ccnA* |
| Spn004 | GGCCCAAATAAGAGCACATGATCC | | |
| **For construction of strain NRD10069 (Δ*ccnB::*$P_c$-[kanR-rpsL$^+$])** | | | |
| Spn005 | ATCATAGACAAGGCGACTGGTAAGGC | IU1945 | Upstream of *ccnB* |
| Spn006 | CCATTAAAAATCAAACGGATCCTAGGAGGTCTTTATTTAATAACTACATG | | |
| kanrpsL For | TAGGATCCGTTTGATTTTTAATGGATAATG | K272 | $P_c$-[*kan rpsL*$^+$][b] |
| kanrpsL rev | GGGCCCCTTTCCTTATGCTTTTG | | |
| Spn007 | TCCAAAAGCATAAGGAAAGGGGCCCCAGGTGGAGTTTTTTAGCTCTATTTCAG | IU1945 | Downstream of *ccnB* |
| Spn008 | CCAAATGAACACTACGACTACCCTCACC | | |
| **For construction of strains NRD10070 and NRD10078 (Δ*ccnC::*$P_c$-[kanR-rpsL$^+$])** | | | |
| Spn036 | GCCTTATCATATCGAGTTGGATCGCTTGC | IU1945 | Upstream of *ccnC* |
| Spn034 | CACATTATCCATTAAAAATCAAACGGATCCTAGTCTATAGTATACCCGACCTATCTTAAAC | | |
| kanrpsL For | TAGGATCCGTTTGATTTTTAATGGATAATG | K272 | $P_c$-[*kan rpsL*$^+$][b] |
| kanrpsL rev | GGGCCCCTTTCCTTATGCTTTTG | | |
| Spn035 | CGTCCAAAAGCATAAGGAAAGGGGCCCGTGTTGGGATTCATGATATAATAATAAAATCG | IU1945 | Downstream of *ccnC* |
| Spn037 | TAATCCCCATCAATGACCCCAACTGAGT | | |
| **For construction of strains NRD10071 and NRD10080 (Δ*ccnD::*$P_c$-[kanR-rpsL$^+$])** | | | |
| **Spn020** | AATGAGTTAGAGCCTGGGGATGTCC | IU1945 | Upstream of *ccnD* |
| **Spn018** | ATCCATTAAAAATCAAACGGATCCTAGAACTTAGTGTACACTCCCTAGCTTAAAGTTTCC | | |
| kanrpsL For | TAGGATCCGTTTGATTTTTAATGGATAATG | K272 | $P_c$-[*kan rpsL*$^+$][b] |

| | | | |
|---|---|---|---|
| kanrpsL rev | GGGCCCCTTTCCTTATGCTTTTG | | |
| Spn019 | CGTCCAAAAGCATAAGGAAAGGGG CCCGAAAAATGGGCTTGGTGCCTGA GAAT | IU1945 | Downstream of *ccnD* |
| Spn021 | GTCAGGTAATTCTCCAAGGGAATGG | | |
| **For construction of strains NRD10072 and NRD10083 (Δ*ccnE::*P$_c$-[kanR-rpsL$^+$])** | | | |
| Spn023 | GTATCGGTGACACCTATTTCTCTGA | IU1945 | Upstream of *ccnE* |
| Spn022 | CACATTATCCATTAAAAATCAAACGG ATCCTAGATTATAGTATACACATCTA ATCTT | | |
| kanrpsL For | TAGGATCCGTTTGATTTTTAATGGAT AATG | K272 | P$_c$-[*kan rpsL*$^+$][b] |
| kanrpsL rev | GGGCCCCTTTCCTTATGCTTTTG | | |
| Spn024 | CGTCCAAAAGCATAAGGAAAGGGG CCCTCATTCATGATATAATAGAAGCA AACGGAG | IU1945 | Downstream of *ccnE* |
| Spn025 | CACTGATTTCAGAACTCTCTACTTCT ATCC | | |
| **For construction of strain NRD10073 (Δ*ccnA*)** | | | |
| Spn013 | CAGTGGAAAAGCATGCGGAGGATTT G | IU1945 | Upstream of *ccnA* |
| Spn015 | TCTATCACCAGCACCTCACCACGCG ACTATATAATACTAGACCATCCT | | |
| Spn014 | AGGATGGTCTAGTATTATATAGTCG CGTGGTGAGGTGCTGGTGATAGA | IU1945 | Downstream of *ccnA* |
| Spn004 | GGCCCAAATAAGAGCACATGATCC | | |
| **For construction of strain NRD10074 (Δ*ccnB*)** | | | |
| Spn013 | CAGTGGAAAAGCATGCGGAGGATTT G | IU1945 | Upstream of *ccnB* |
| Spn028 | GTCCCCAAAAGCCTGAAATAGAGCT AACTACATGATACAAGACGAAACTTA AAAC | | |
| Spn029 | AAGTTTCGTCTTGTATCATGTAGTTA GCTCTATTTCAGGCTTTTGGGGACT ATTC | IU1945 | Downstream of *ccnB* |
| Spn004 | GGCCCAAATAAGAGCACATGATCC | | |
| **For construction of strains NRD10075 and NRD10079 (Δ*ccnC*)** | | | |
| Spn036 | GCCTTATCATATCGAGTTGGATCGC TTGC | IU1945 | Upstream of *ccnC* |
| 5-ccnCcleanKO Rev | CGATTTTATTATTATATCATGAATCC CAACACGTCTATAGTATACCCGACC TATCTTAAAC | | |
| 3'ccnCcleanKO For | GTTTAAGATAGGTCGGGTATACTAT AGACGTGTTGGGATTCATGATATAA TAATAAAATCG | IU1945 | Downstream of *ccnC* |
| Spn037 | TAATCCCCATCAATGACCCCAACTG AGT | | |
| **For construction of strains NRD10076 and NRD10081 (Δ*ccnD*)** | | | |
| Spn020 | AATGAGTTAGAGCCTGGGGATGTCC | IU1945 | |

| | | | |
|---|---|---|---|
| Spn031 | TTCTCAGGCACCAAGCCCATTTTTCGAACTTAGTGTACACTCCCTAGCTT | | Upstream of *ccnD* |
| Spn030 | CTTTAAGCTAGGGAGTGTACACTAAGTTCGAAAAATGGGCTTGGTGCCTGAGAAT | IU1945 | Downstream of *ccnD* |
| Spn021 | GTCAGGTAATTCTCCAAGGGAATGG | | |
| **For construction of strains NRD10077 and NRD10085 (Δ*ccnE*)** | | | |
| Spn023 | GTATCGGTGACACCTATTTCTCTGA | IU1945 | Upstream of *ccnE* |
| Spn033 | CCGTCCTCCGTTTGCTTCTATTATATCATGAATGAGATTATAGTATACACATCTAATCTT | | |
| Spn032 | AAGATTAGATGTGTATACTATAATCTCATTCATGATATAATAGAAGCAAACGGAGGACGG | IU1945 | Downstream of *ccnE* |
| Spn025 | CACTGATTTCAGAACTCTCTACTTCTATCC | | |
| **For construction of strain NRD10171 (Δ*ccnAB::*$P_c$-[kanR-rpsL$^{+}$])** | | | |
| Spn001 | TCATAGACAAGGCGACTGGTAAGG | IU1945 | Upstream of *ccnA* |
| Spn002 | CCATTAAAAATCAAACGGATCCTAAAAAAAGTTTAGGATTTTATTAAATAAAGTTAGG | | |
| kanrpsL For | TAGGATCCGTTTGATTTTTAATGGATAATG | K272 | $P_c$-[*kan rpsL*$^{+}$][b] |
| kanrpsL rev | GGGCCCCTTTCCTTATGCTTTTG | | |
| Spn007 | TCCAAAAGCATAAGGAAAGGGGCCCCAGGTGGAGTTTTTTAGCTCTATTTCAG | IU1945 | Downstream of *ccnB* |
| Spn008 | CCAAATGAACACTACGACTACCCTCACC | | |
| **For construction of strain NRD10176 (Δ*ccnAB*)** | | | |
| Spn001 | TCATAGACAAGGCGACTGGTAAGG | IU1945 | Upstream of *ccnA* |
| 53-ccnAB For2 | GCAAAATTTAAGGATGGTCTAGTATTATATAGTCAGCTCTATTTCAGGCTTTTGGG | | |
| 53-ccnAB Rev2 | CCCAAAAGCCTGAAATAGAGCTGACTATATAATACTAGACCATCCTTAAATTTTGC | IU1945 | Downstream of *ccnB* |
| Spn008 | CCAAATGAACACTACGACTACCCTCACC | | |
| **For construction of strains NRD10225 and NRD10306 (Δ*ccnA::*$P_c$-[kanR-rpsL$^{+}$])** | | | |
| T4ccnA For | CAGTGGAAAAGCATGCGGAGG | TIGR4 | Upstream of *ccnA* |
| 5-T4ccnA-rpsL Rev | CATTATCCATTAAAAATCAAACGGATCCTAGACTATATGATACTAGACCATCCTTAAAC | | |
| kanrpsL For | TAGGATCCGTTTGATTTTTAATGGATAATG | K272 | $P_c$-[*kan rpsL*$^{+}$][b] |
| kanrpsL rev | GGGCCCCTTTCCTTATGCTTTTG | | |
| 3-T4ccnA-rpsL For | CAAAAGCATAAGGAAAGGGGCCCGGTACGACGGGCATGTCG | TIGR4 | Downstream of *ccnA* |

| | | | |
|---|---|---|---|
| T4ccnA Rev | GGCTCATGACTTGGACAATGG | | |
| **For construction of strains NRD10230 and NRD10312 (Δ*ccnA*)** | | | |
| T4ccnA For | CAGTGGAAAAGCATGCGGAGG | TIGR4 | Upstream of *ccnA* |
| 5-T4ccnA-cln Rev | CGACATGCCCGTCGTACCGCAGACT ATATGATACTAGACCATCCTTAAAC | | |
| 3-T4ccnA-cln-For | GTTTAAGGATGGTCTAGTATCATATA GTCTGCGGTACGACGGCATGTCG | TIGR4 | Downstream of *ccnA* |
| T4ccnA Rev | GGCTCATGACTTGGACAATGG | | |
| **For construction of strain NRD10247 (Δ*ccnB::*$P_c$-[kanR-rpsL$^+$])** | | | |
| T4ccnB For | GCCATTGTCCAAGTCATGAGC | TIGR4 | Upstream of *ccnB* |
| 5-T4ccnB-rpsL Rev | CATTATCCATTAAAAATCAAACGGAT CCTACTACATGATACAAGACGAAAC TTAAAACTAGC | | |
| kanrpsL For | TAGGATCCGTTTGATTTTTAATGGAT AATG | K272 | $P_c$-[*kan rpsL*$^+$][b] |
| kanrpsL rev | GGGCCCCTTTCCTTATGCTTTTG | | |
| 3-T4ccnB-rpsL For | CAAAAGCATAAGGAAAGGGGCCCA GCTCTATTTCAGGATTTTTGGGAC | TIGR4 | Downstream of *ccnB* |
| T4ccnB Rev | GAGGACATCTCCAGGCTCTAAATC | | |
| **For construction of strains NRD10249 and NRD10322 (Δ*ccnB*)** | | | |
| T4ccnB For | GCCATTGTCCAAGTCATGAGC | TIGR4 | Upstream of *ccnB* |
| 5-T4ccnB-cln Rev | GTCCCAAAAATCCTGAAATAGAGCT AACTACATGATACAAGACGAAACTTA AAACTAGC | | |
| 3-T4ccnB-cln-For | GCTAGTTTTAAGTTTCGTCTTGTATC ATGTAGTTAGCTCTATTTCAGGATTT TTGGGAC | TIGR4 | Downstream of *ccnB* |
| T4ccnB Rev | GAGGACATCTCCAGGCTCTAAATC | | |
| **For construction of strains NRD10251 and NRD10330 (Δ*ccnC::*$P_c$-[kanR-rpsL$^+$])** | | | |
| T4ccnC For | GCCTTATCATATCGAGTTGGATCGC TTGC | TIGR4 | Upstream of *ccnC* |
| 5-T4ccnC-rpsL Rev | CATTATCCATTAAAAATCAAACGGAT CCTAGTCTATAGTATACCCGACCTAT CTTAAAC | | |
| kanrpsL For | TAGGATCCGTTTGATTTTTAATGGAT AATG | K272 | $P_c$-[*kan rpsL*$^+$][b] |
| kanrpsL rev | GGGCCCCTTTCCTTATGCTTTTG | | |
| 3-cT4cnC-rpsL For | CAAAAGCATAAGGAAAGGGGCCCG TGTTGGGATTCATGATATAATAATAA AATCG | TIGR4 | Downstream of *ccnC* |
| T4ccnC Rev | CTTCAAGAACAACCTCTTGGTCTG | | |
| **For construction of strains NRD10254 and NRD10332 (Δ*ccnC*)** | | | |
| T4ccnC For | GCCTTATCATATCGAGTTGGATCGC TTGC | TIGR4 | Upstream of *ccnC* |
| 5-T4ccnC-cln Rev | CGATTTTATTATTATATCATGAATCC CAACACGTCTATAGTATACCCGACC TATCTTAAAC | | |

| | | | |
|---|---|---|---|
| 3-T4ccnC-cln-For | GTTTAAGATAGGTCGGGTATACTATAGACGTGTTGGGATTCATGATATAATAATAAAATCG | TIGR4 | Downstream of *ccnC* |
| T4ccnC Rev | CTTCAAGAACAACCTCTTGGTCTG | | |
| **For construction of strains NRD10257 and NRD10336 (Δ*ccnD::*$P_c$-[kanR-rpsL$^+$])** | | | |
| T4ccnD For | AATGATTTAGAGCCTGGAGATGTCC | TIGR4 | Upstream of *ccnD* |
| 5-T4ccnD-rpsL Rev | CATTATCCATTAAAAATCAAACGGATCCTAGAACTTAGTGTACACTCCCTAGCTTAAAG | | |
| kanrpsL For | TAGGATCCGTTTGATTTTTAATGGATAATG | K272 | $P_c$-[*kan rpsL*$^+$][b] |
| kanrpsL rev | GGGCCCCTTTCCTTATGCTTTTG | | |
| 3-T4ccnD-rpsL For | CAAAAGCATAAGGAAAGGGGCCCGAAAAATGGGCTTGGTGCCTG | TIGR4 | Downstream of *ccnD* |
| T4ccnD Rev | GTCAGGTAATTCTCCAAGGGAATGG | | |
| **For construction of strains NRD10261 and NRD10340 (Δ*ccnD*)** | | | |
| T4ccnD For | AATGATTTAGAGCCTGGAGATGTCC | TIGR4 | Upstream of *ccnD* |
| 5-T4ccnD-cln Rev | CAGGCACCAAGCCCATTTTTCGAACTTAGTGTACACTCCCTAGCTTAAAG | | |
| 3-T4ccnD-cln-For | CTTTAAGCTAGGGAGTGTACACTAAGTTCGAAAAATGGGCTTGGTGCCTG | TIGR4 | Downstream of *ccnD* |
| T4ccnD Rev | GTCAGGTAATTCTCCAAGGGAATGG | | |
| **For construction of strains NRD10265 and NRD10344 (Δ*ccnE::*$P_c$-[kanR-rpsL$^+$])** | | | |
| T4ccnE For | GTATCGGTGACACCTATTTCTCTGA | TIGR4 | Upstream of *ccnE* |
| 5-T4ccnE-rpsL Rev | CATTATCCATTAAAAATCAAACGGATCCTAGATTATAGTATACACGTCTAATCTTAAATAGAAC | | |
| kanrpsL For | TAGGATCCGTTTGATTTTTAATGGATAATG | K272 | $P_c$-[*kan rpsL*$^+$][b] |
| kanrpsL rev | GGGCCCCTTTCCTTATGCTTTTG | | |
| 3-T4ccnE-rpsL For | CAAAAGCATAAGGAAAGGGGCCCTCATTCATGATATAATAGAAGCAAACGG | TIGR4 | Downstream of *ccnE* |
| T4ccnE Rev | GCTATCTGGTATCATTTCTGGAGCC | | |
| **For construction of strains NRD10266 and NRD10345 (Δ*ccnE*)** | | | |
| T4ccnE For | GTATCGGTGACACCTATTTCTCTGA | TIGR4 | Upstream of *ccnE* |
| 5-T4ccnE-cln Rev | CCGTTTGCTTCTATTATATCATGAATGAGATTATAGTATACACGTCTAATCTTAAATAGAAC | | |
| 3-T4ccnE-cln-For | GTTCTATTTAAGATTAGACGTGTATACTATAATCTCATTCATGATATAATAGAAGCAAACGG | TIGR4 | Downstream of *ccnE* |
| T4ccnE Rev | GCTATCTGGTATCATTTCTGGAGCC | | |
| **For construction of strains NRD10311 and NRD10346 (*rpsL*$^+$*-rpsG*$^+$*-cat*)** | | | |
| KK265 | GTGAATTGGTCGGAATTGTAGCTAACAG | TIGR4 | Upstream of *rpsL* + stop codon of *rpsG* |
| KK529 | GCCTCCTAAATTTACCAACGGAAGTGTGCGAATG | | |

| | | | |
|---|---|---|---|
| KK530 | CCGTTGGTAAATTTAGGAGGCATAT CAAATGAACT | IU9681 | *cat* cassette |
| KK531 | CGCATCCTATCTTATAAAAGCCAGT CATTAGGCC | | |
| KK532 | CTGGCTTTTATAAGATAGGATGCGA AAGCGTTAAG | TIGR4 | Stop codon of *rpsG* + downstream of *rpsG* |
| KK270 | CAAGGATATCCGTACCAAGGTCGTT AG | | |
| **For construction of strains NRD10390 and NRD10391 (*CEP::*$P_c$-[kanR-rpsL$^+$])** | | | |
| AmiF For | CCAGCTGTGATTGCAAATCTCCATG | IU5122 | Upstream of *amiF* + downstream of *amiF* |
| TreR Rev | GGCTTCTTGTTCAAATTTTCCCATTT GATTCTC | | |
| **For construction of strain NRD10393 (*CEP::*$T_1$-$T_2$-*ccnA-ccnB*)** | | | |
| 5-AmiF-Ptet for | CTGGCTGACTAGGAGGAAGG | IU5382 | Upstream of *amiF* + $T_1$-$T_2$ terminator |
| 5-ccnABCEP_rev | CCAAGTCATCCCTATACAATTATAGG TGGGTAGAAACGCAAAAAGGCCATC | | |
| M-ccnAB For | GATGGCCTTTTTGCGTTTCTACCCA CCTATAATTGTATAGGGATGACTTG G | IU1945 | 101 nt upstream of *ccnA* + 139 nt downstream of *ccnB* |
| M-ccnAB Rev | GCTCCCTTTTTTAATGGTAACACCG CTGAAGCGACCACAGACC | | |
| 3-ccnABCEP_ for | AGGTCTGTGGTCGCTTCAGCGGTGT TACCATTAAAAAAGGGAGC | IU5382 | 26 nt of spd_1082 + downstream of *amiF* |
| 3-TreR-Ptet rev | CCCATTTGATTCTCCTTATACTTGTC AAAGC | | |
| **For construction of strain NRD10394 (*CEP::*$T_1$-$T_2$-*ccnC*)** | | | |
| 5-AmiF-Ptet for | CTGGCTGACTAGGAGGAAGG | IU5382 | Upstream of *amiF* + $T_1$-$T_2$ terminator |
| 5-ccnCCEP_rev | CGAACGCCAACCAATTCACTCGTAG AAACGCAAAAAGGCCATC | | |
| M-ccnC For | GATGGCCTTTTTGCGTTTCTACGAG TGAATTGGTTGGCGTTCG | IU1945 | 133 nt upstream of *ccnC* + 111 nt downstream of *ccnC* |
| M-ccnC Rev | GCTCCCTTTTTTAATGGTAACACCG CCTCTCTCAAAGCCTCCC | | |
| 3-ccnCCEP_ for | GGGAGGCTTTGAGAGAGGCGGTGT TACCATTAAAAAAGGGAGC | IU5382 | 26 nt of spd_1082 + downstream of *amiF* |
| 3-TreR-Ptet rev | CCCATTTGATTCTCCTTATACTTGTC AAAGC | | |
| **For construction of strains NRD10396 and NRD10397 (Δ*bgaA-kan*-$T_1$-$T_2$-*ccnD*)** | | | |
| 5-BgaFor2 | CTGGTGATATCAAAGCAATCCTTGG | IU10508 | Upstream of bgaA + $T_1$-$T_2$ terminator |
| 5-ccnDBga Rev | CCATTTCTCCTGCCAATTTTTCTTGG TAGAAACGCAAAAAGGCCATC | | |
| M-ccnD For | GATGGCCTTTTTGCGTTTCTACCAA GAAAAATTGGCAGGAGAAATGG | IU1945 | 86 nt upstream of *ccnD* + 55 nt downstream of *ccnD* |
| M-ccnD Rev | GCAACTGGTTTATGAGAAAGTAAGT TCCCCCATTTTCTTCTATCACTAAGC | | |
| 3-ccnDBga For | GCTTAGTGATAGAAGAAAATGGGGG AACTTACTTTCTCATAAACCAGTTGC | IU10508 | 1501 nt of *bgaA* + |

| | | | |
|---|---|---|---|
| 3-Bga-rpsLKan Rev | CTGGTTTTTCCTTAGTCAACTGGATACGG | | downstream of *bgaA* |
| **For construction of strains NRD10441 and NRD10443 (Δ*psaR::*$P_c$-[kanR-rpsL$^+$])** | | | |
| 5-psaR for | GAGGCTACCCTGCCTCTACTC | IU1945 | Upstream of *psaR* |
| 5-psaR rpsL rev | CATTATCCATTAAAAATCAAACGGATCCTATAGATAGTCTTCTTTGTTTGGGGTC | | |
| kanrpsL For | TAGGATCCGTTTGATTTTTAATGGATAATG | K272 | $P_c$-[*kan rpsL*$^+$][b] |
| kanrpsL rev | GGGCCCCTTTCCTTATGCTTTTG | | |
| 3-psaR rpsL for | CAAAAGCATAAGGAAAGGGGCCCATTGCAAAACAACTCTATGTCGAG | IU1945 | Downstream of *psaR* |
| 3-psaR rev | CCAGAGAGCAAGAGCCACTC | | |
| **For construction of strains NRD10442 and NRD10444 (Δ*mntE::*$P_c$-[kanR-rpsL$^+$])** | | | |
| 5-mntE for | CCGCATCTTGAAGCATACCAGC | IU1945 | Upstream of *mntE* |
| 5-mntE rpsL rev | CATTATCCATTAAAAATCAAACGGATCCTACTCAGCTAACTTGAGATTTGAGATAG | | |
| kanrpsL For | TAGGATCCGTTTGATTTTTAATGGATAATG | K272 | $P_c$-[*kan rpsL*$^+$][b] |
| kanrpsL rev | GGGCCCCTTTCCTTATGCTTTTG | | |
| 3-mntE rpsL for | CAAAAGCATAAGGAAAGGGGCCCTGGCAAAATATCTTTCATCAAGAAACC | IU1945 | Downstream of *mntE* |
| 3-mntE rev | GGAAGAAAGTCATCGAGTTTCAGG | | |
| **For construction of strains NRD10447 and NRD10449 (Δ*psaR*)** | | | |
| 5-psaR for | GAGGCTACCCTGCCTCTACTC | IU1945 | Upstream of *psaR* |
| 5-psaR cln rev | CTCGACATAGAGTTGTTTTGCAATTAGATAGTCTTCTTTGTTTGGGGTC | | |
| 3-psaR cln for | GACCCCAAACAAAGAAGACTATCTAATTGCAAAACAACTCTATGTCGAG | IU1945 | Downstream of *psaR* |
| 3-psaR rev | CCAGAGAGCAAGAGCCACTC | | |
| **For construction of strains NRD10448 and NRD10450 (Δ*mntE*)** | | | |
| 5-mntE for | CCGCATCTTGAAGCATACCAGC | IU1945 | Upstream of *mntE* |
| 5-mntE cln rev | GGTTTCTTGATGAAAGATATTTTGCCACTCAGCTAACTTGAGATTTGAGATAG | | |
| 3-mntE cln for | CTATCTCAAATCTCAAGTTAGCTGAGTGGCAAAATATCTTTCATCAAGAAACC | IU1945 | Downstream of *mntE* |
| 3-mntE rev | GGAAGAAAGTCATCGAGTTTCAGG | | |
| **For construction of strains NRD10533 and NRD10534 (Δ*sodA::erm*)** | | | |
| 5-sodA For | CAGGGTCAGTTTGAAGTGATGAAGAG | IU1945 | Upstream of *sodA* |
| 5-sodA-erm Rev | TATTTTATATTTTTGTTCATCTGTAATACCTCTTTTTCTTTCTATATG | | |
| M-sodA-erm For | GAAAAAGAGGTATTACAGATGAACAAAAATATAAAATATTCTC | NRD10491[c] | erm[b] |
| M-sodA-erm Rev | CCTCCAACTATCATTATTTCCTCCCGTTAAATAATAGATAAC | | |

| | | | |
|---|---|---|---|
| 3-sodA-erm For | TAACGGGAGGAAATAATGATAGTTGGAGGGAAGAATTGTTC | IU1945 | stop codon of *sodA* + downstream |
| 3-sodA Rev | CCATAGTGTTGACGCATGAGTACAG | | |
| **For construction of strains TIGR4S and TIGR4SΔcps (*rpsLK56T*)** | | | |
| HE01 | GCCGTAGTCATCTTTCTTGGCATC | IU11966 | Upstream *rpsL* + 181 nt of *rpsL* with A167C change |
| HE02 | CTGAGTTAGGTTTTGTAGGTGTCATTGTTC | | |
| HE03 | GAACAATGACACCTACAAAACCTAACTCAG | IU11966 | 263 nt of *rpsL* with A167C change + downstream |
| HE04 | CTAATTTGAACCCGGGCTAAAGTTAG | | |

[a] Genomic DNA of indicated *S. pneumoniae* strains was used as templates for PCR reactions, except for $P_c$-[*kan-rpsL*$^+$] and $P_c$-*erm* cassettes.

[b] $P_c$-*erm* and $P_c$-[*kan-rpsL*$^+$] cassettes are described in (Tsui *et al.*, 2010) (*8*).

[c] NRD10491 is an unpublished strain with genotype D39 Δcps pnp-3XFLAG-$P_c$-*erm*.

**Table S6.** Oligonucleotide primers and probes used for qRT-PCR and northern blots.

| **Primer** | **Primer or probe sequence (5'-3')** | **Gene** |
|---|---|---|
| D39_czcD_For | CGGGCTCTGTTCTAGTCATTT | *czcD* |
| D39_czcD_Rev | CCAGACTCGCTAACAGATTGAT | *czcD* |
| TIGR4_czcD_For | TGGTGGTTGGTAAGGGAAAG | *czcD* |
| TIGR4_czcD_Rev | AGAACAATCGCCATCAGGATAA | *czcD* |
| piuB_For | CGGCTCAGTCACAGAAGTTATC | *piuB* |
| piuB_Rev | GCCTAAGAAGAGCCACTCATAC | *piuB* |
| spd_1267_For | TGGATGAGCCTTCATCGAATTTA | *spd_1267* |
| spd_1267_Rev | CACGGTCAACTATGTCCATCAA | *spd_1267* |
| tuf_For | ATCACTGGTGCTGCTCAAA | *tuf* |
| tuf_Rev | CCTGACGTGAAAGAAGGATGT | *tuf* |
| **Probe** | | |
| sodA | GCAAGCAAGGCTTCAAGGTCTTCACCGATTTCAGG | *sodA* |
| 5s rRNA | GCGTTCTAGGGCTTAACTTCTGTGTTCGGCATGGG | *rrfA* |

**SUPPLEMENTAL FIGURES**

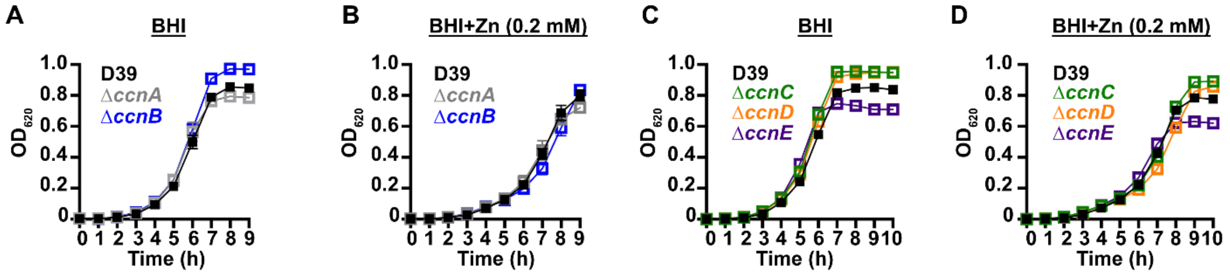


**FIGURE S1.** Growth phenotypes of *S. pneumoniae* D39 derived strains harboring deletion of individual *ccn* genes. Growth characteristics at 37°C under an atmosphere of 5% $CO_2$ in BHI broth alone (A,C) or with 0.2 mM $ZnSO_4$ (B, D) of the following strains: (A, B) IU781 (D39), NRD10073 (Δ*ccnA*), and NRD10074 (Δ*ccnB*); (C, D) IU781 (D39), NRD10075 (Δ*ccnC*), NRD10076 (Δ*ccnD*), and NRD10077 (Δ*ccnE*). Each point on the graph represents the mean $OD_{620}$ value from three independent cultures. Error bars, which in some cases are too small to observe in the graph, represent the standard deviation (SD).

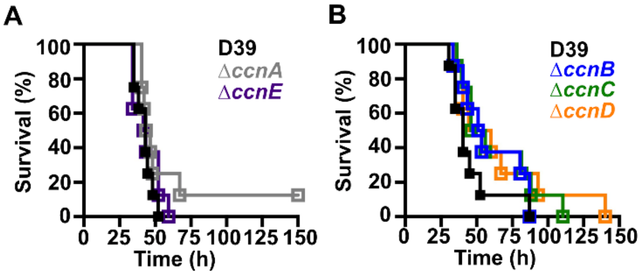


**FIGURE S2.** Virulence phenotypes of *S. pneumoniae* strains harboring deletion of individual *ccn* genes. Survival curve of ICR outbred mice after infection with ~$10^7$ CFU in a 50 μL inoculum of the following *S. pneumoniae* strains: (A) IU781 (D39), NRD10073 (Δ*ccnA*), and NRD10077 (Δ*ccnE*); (B) IU781 (D39), NRD10074 (Δ*ccnB*), NRD10075 (Δ*ccnC*), and NRD10076 (Δ*ccnD*). Eight mice were infected per strain. Disease progression of animals was monitored, the time at which animals reached a moribund state was recorded, and these mice were subsequently euthanized as described in Materials and Methods. A survival curve was generated from this data and analyzed by Kaplan-Meier statistics and log rank test to determine P-values.

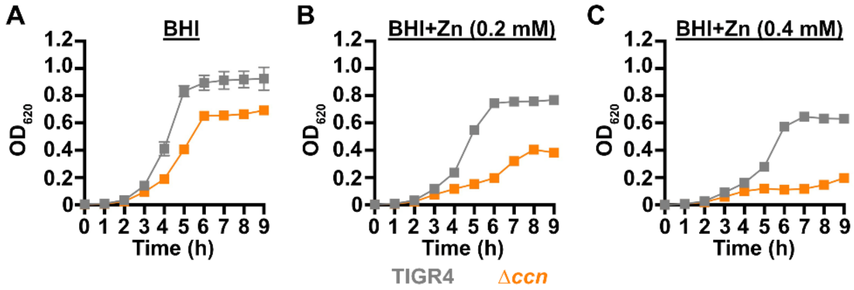


**FIGURE S3.** Growth phenotypes of *S. pneumoniae* TIGR4 derived strains harboring deletion of the *ccn* genes. Growth characteristics at 37°C under an atmosphere of 5% $CO_2$ in BHI broth alone (A) or with 0.2 mM (B) or 0.4 mM (C) $ZnSO_4$ of NRD10311 (TIGR4; TIGR4 *rpsL*$^{+}$-*rpsG*$^{+}$-*cat*) and NRD10346 (Δ*ccn*; TIGR4 *rpsL*$^{+}$-*rpsG*$^{+}$-*cat* Δ*ccnABCDE*). Each point on the graph represents the mean $OD_{620}$ value from three independent cultures. Error bars, which in some cases are too small to observe in the graph, represent the standard deviation (SD).

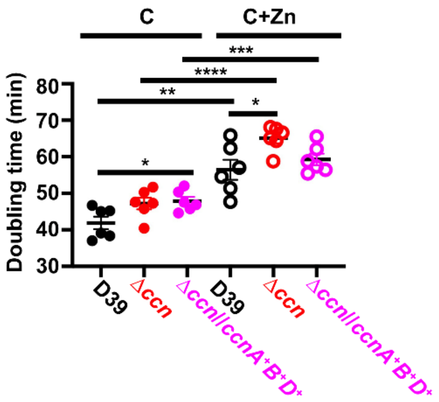


**FIGURE S4.** Doubling times of *S. pneumoniae* D39, Δ*ccnABCDE* mutant, or Δ*ccnABCDE* mutant strain complemented with *ccnA, ccnB,* and *ccnC* in C medium alone or supplemented with Zn. Shown are the mean doubling times during exponential growth of IU1781 (D39), NRD10176 (Δ*ccn)*, and NRD10396 (Δ*ccn//ccnA*$^+$*B*$^+$*D*$^+$) grown in C medium alone or supplemented with 0.2 mM ZnS04 as described in *Materials and Methods*. Doubling times for individual replicates are shown with solid lines indicating the mean of six different biological replicates and error bars denoting standard error of the mean (SEM). Statistical significance as determined by a Mann-Whitney test is indicate as * (P < 0.05), ** (P < 0.005), *** (P < 0.0005), or **** (P < 0.00005).

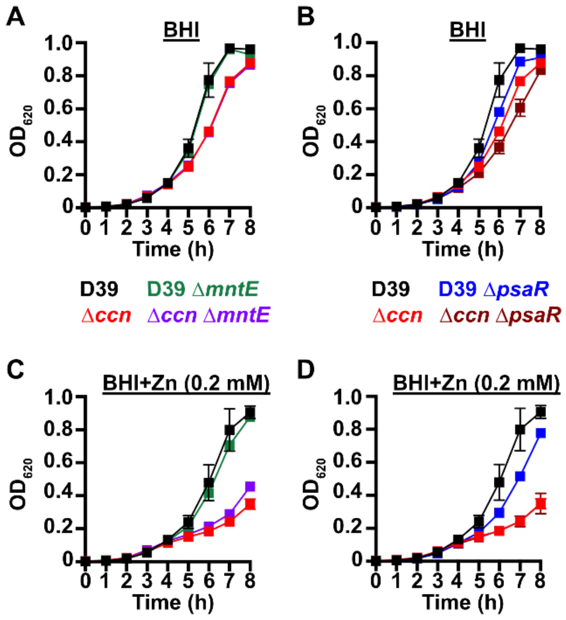


**FIGURE S5.** Growth phenotypes of *S. pneumoniae* D39 and derived strains harboring deletion of the *ccn* genes and/or *psaR* or *mntE*. Growth characteristics at 37°C under an atmosphere of 5% $CO_2$ in BHI broth alone (A, B) or with 0.2 mM $ZnSO_4$ (C, D) of the following strains: (A, C) IU781 (D39), NRD10176 (Δ*ccnABCDE*), NRD10448 (Δ*mntE*), and NRD10450 (Δ*ccnABCDE* Δ*mntE*); (B, D) IU781 (D39), NRD10176 (Δ*ccnABCDE*), NRD10447 (Δ*psaR*), and NRD10450 (Δ*ccnABCDE* Δ*psaR*). Each point on the graph represents the mean $OD_{620}$ value from three independent cultures. Error bars, which in some cases are too small to observe in the graph, represent the standard deviation (SD).
